# 4-Thiaproline accelerates the slow folding phase of proteins containing *cis* prolines in the native state by two orders of magnitude

**DOI:** 10.1101/2023.06.23.546227

**Authors:** Jennie O’ Loughlin, Kirill Zinovjev, Silvia Napolitano, Marc van der Kamp, Marina Rubini

## Abstract

The *cis/trans* isomerization of peptidyl-prolyl peptide bonds is often the bottleneck of the refolding reaction for proteins containing *cis* proline residues in the native state. Proline (Pro) analogues, especially C4-substituted fluoroprolines, have been widely used in protein engineering to enhance the thermodynamic stability of peptides and proteins and to investigate folding kinetics. 4-thiaproline (Thp) has been shown to bias the ring pucker of Pro, to increase the *cis* population percentage of model peptides in comparison to Pro, and to diminish the activation energy barrier for the *cis/trans* isomerization reaction. Despite its intriguing properties, Thp has been seldom incorporated into proteins. Moreover, the impact of Thp on the folding kinetics of globular proteins has never been reported. In this study, we show that upon incorporation of Thp at *cis*Pro76 into the thioredoxin variant Trx1P the half-life of the refolding reaction decreased from ∼2 hours to ∼35 seconds. A dramatic acceleration of the refolding rate could be observed also for the protein pseudo wild-type barstar upon replacement of *cis*Pro48 with Thp. Quantum chemical calculations revealed that the replacement of the C^γ^H_2_ group by a sulfur atom in the pyrrolidine ring, lowers the barrier for *cis*/*trans* rotation due to a weakened peptide bond. The protein variants retained their thermodynamic stability upon incorporation of Thp, while the catalytic and enzymatic activities of the modified Trx1P remained unchanged. Our results show that the Pro isostere Thp might eliminate the bottleneck of the refolding reaction of proteins containing *cis* proline residues in the native state, independent from the local structural environment.

## Introduction

The *cis/trans* isomerization of peptidyl-prolyl peptide bonds is an intrinsically slow reaction that significantly curbs the folding process in proteins that contain *cis* proline (Pro) residues in the native state.^1-2^ In living organisms, peptidyl-prolyl *cis/trans* isomerases (PPIases) catalyse this rate-limiting folding step, by diminishing the energy barrier between the *trans* and *cis* peptide bond conformation, thereby accelerating the folding process from minutes to milliseconds.^3^ In nature, the switching between these two conformational states is associated with modulation of protein function and the regulation of cellular processes.^4^ Therefore, it is not surprising that PPIases play a pivotal role in health and disease.^5-6^ It has been shown that *trans* to *cis* isomerization of Pro residues can be the bottleneck also for *in vitro* refolding reactions for proteins containing prolines in the *cis* conformation in their native three-dimensional structure.^7^

Pro displays unique features in comparison to the other proteinogenic amino acids. The pyrrolidine ring confers an exceptional conformational constraint on the phi angle in the peptide bond, due to the constriction of the Cα and nitrogen atoms of Pro in the ring.^8^ While amino acids other than Pro adopt almost always the *trans* conformation of the peptide bond in order to avoid steric repulsion between the Cα atom and the Cα atom of the preceding residue, the *cis* and *trans* conformations of peptidyl-prolyl peptide bonds are almost isoenergetic.^9^ In fact, although the *trans* conformation is still preferred in Xaa-Pro bond (Xaa: any amino acid), steric repulsions cannot be fully eliminated,^10^ thus making *cis* prolyl peptide bonds more frequent (∼6 % of all prolines) in the context of proteins tertiary structures.^11^ The pyrrolidine ring adopts mainly two pucker conformations, with the C^γ^ atom either pointing towards the opposite side of the carbonyl group of Pro (*exo* conformation), or being on the same side of the plane (*endo* conformation). The adoption of the *endo* ring pucker is preferred when Pro residues are involved in a *cis* peptide bond, while no pucker preference is shown in the presence of a *trans* peptide bond.^12^

Proline analogues have been extensively used in protein engineering to introduce new chemical functionalities while maintaining conformational constraints.^13^ By installing an electron-withdrawing group such as fluorine on the C^γ^ atom, the pucker preference of the ring can be biased based on the gauche effect.^14^ This in turn, can affect the *cis/trans* equilibrium of the peptide bond.^15^ The *endo* pucker conformation elicits the stability of the *cis* peptide bond, while the *exo* conformation goes along with an increased stability of the *trans* peptide bond. In fact, it has been shown that (4*S*)-fluoroproline ((4*S*)-Flp) promotes the *endo* pucker while increasing the stability of the *cis*-peptide bond, whereas (4*R*)-fluoroproline ((4*R*)-Flp) favours the adoption of *exo* pucker and the stability of the *trans*-peptide bond.^16-17^ For this reason, 4-fluoroproline analogues have been used both to increase the thermodynamic and conformational stability of proteins,^18-23^ and to investigate protein folding kinetics.^24-26^ In fact, the presence of the electron-withdrawing fluorine lowers the energy barrier to *cis/trans* isomerization as it decreases the double bond character of the preceding peptide bond.^20^

At present, the pseudoproline thiaproline, 1,3-thiazolidine-4-carboxylic acid (Thp), has been seldom incorporated into proteins,^27-28^ in spite of its intriguing features. Thp displays beneficial pharmaceutical properties due its anti-aging, anti-cancer, and anti-inflammatory activities and it is a promising tool for drug design.^29-31^ Moreover, Thp has proved to be a substrate for human prolyl 4-hydroxylase,^32^ and it was suggested that it could be exploited to design collagen-related biomaterials. ^33^

Although the thiazolidine ring in Thp displays a slightly different geometry in comparison to the pyrrolidine ring of Pro due to the larger size of the sulfur atom, Thp can be considered isosteric to Pro.^34-36^ X-ray crystallography and *ab initio* calculations have shown that in model peptides Thp displays an *endo* puckering ring preference that correlates with a higher *cis* population percentage in comparison to Pro. NMR studies of the *trans/cis* equilibrium of Ac-Thp-OMe, in water at 25 ^°^C, showed the K_*trans/cis*_ to be 2.8,^35^ considerably lower than for Ac-Pro-OMe (K_*trans/cis*_ = 4.6 under the same conditions).^16^ The correlation between the ring pucker and the *cis/trans* amide population can be ascribed to the favourable n → π* interaction between the Pro amide carbonyl oxygen and the carbonyl carbon of the subsequent amide.^16^ In comparison to Pro, Thp exhibits a lower energy barrier between the *endo* and *exo* puckers, particularly in the *trans* conformation, where the *endo* pucker is more stable than the *exo* pucker by only 0.04 kJ/mol, compared to 1.34 kJ/mol for the Pro analogue, as determined by computational studies of Ac-Xaa-OMe peptides (Xaa is Pro or a Pro analogue).^36^ Early NMR studies on model dipeptides Ala-Xaa-(4-)nitroanilide and tetrapeptides Ala-Gly-Xaa-Phe-(4-)nitroanilide, showed that Thp decreases the free energy of activation for peptide isomerization by 10 kJ/mol in comparison to Pro, while the *cis/trans* isomerisation rate constant was increased up to 100-fold compared to the proline peptide.^37^ To date, no data are reported concerning the effect of Thp on protein folding. We reasoned that if the decrease in free energy of activation for *cis/trans* isomerization observed in model peptides would apply also for globular proteins, the slow refolding phase for *in vitro* refolding reactions for proteins containing prolines in the *cis* conformation in the native state could be considerably accelerated. To prove our hypothesis, we chose the model proteins *E. coli* thioredoxin1P (Trx1P) and pseudo wild-type barstar (pwt-b*) from *B. amiloliquefaciens*, for which the folding pathway follows the formation of a native-like long-lived intermediate with a non-native *trans* Pro peptide bond (**Figure 1**).

**Figure 1.**
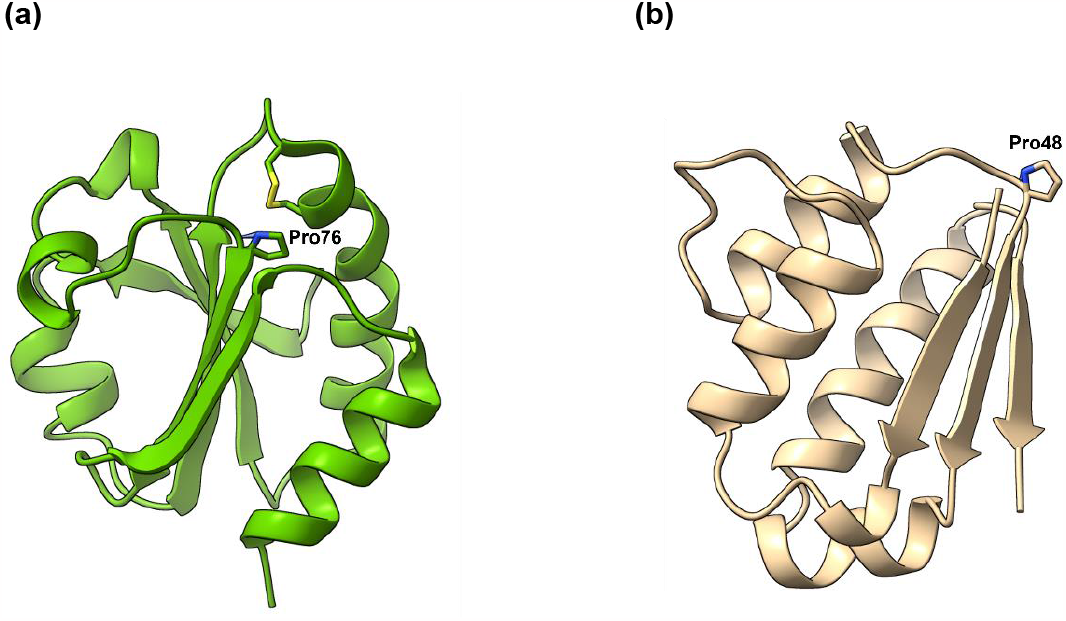
(a) Three-dimensional structure of *E. coli* thioredoxin (PDB: 2TRX). The conserved *cis*Pro76 next to the catalytic disulfide bond is shown (stick representation). (b) Three-dimensional structure of barstar (PDB: 1AY7). The solvent exposed *cis*Pro48 is shown (stick representation).

## Results and Discussion

At first, we tested the effect of Thp on protein folding, thermodynamic stability, and biological activity, by using our well-characterised model protein thioredoxin 1P (Trx1P). Trx1P (Pro34Ala, Pro40Ala, Pro64Ala, Pro68Ala) is an *E. coli* thioredoxin (Trx) variant which contains *cis*Pro76 as the only proline residue whereas the *trans* Pro residues were mutated to Ala.^18^ This conserved *cis* prolyl peptide bond is located in the protein interior next to the catalytic disulfide (Cys32 and Cys35) of the oxidoreductase and is essential for Trx bioactivity and thermodynamic stability.^38^

Thp was introduced into Trx1P at *cis*Pro76 by selective pressure incorporation. Briefly, the proline auxotrophic cell strain CAG18515 transformed with an additional plasmid for overexpression of an engineered *E. coli* prolyl-tRNA-synthetase (C443G) to increase the incorporation of the Pro analogue, was grown in M9 minimal medium.^39^ The best incorporation results were obtained by harvesting the cells 3 h after induction with IPTG (0.5 mM) at 37 ^°^C in the presence of 8 mM Thp. The variant protein Trx1Thp was purified by ion-exchange chromatography and recombinant protein yields were ∼2 mg/L of bacterial culture. ESI-MS analysis showed a Thp incorporation level of ∼85% (Supplementary Figure 1). Previous incorporation experiments reported an almost quantitative replacement of Pro by Thp for the protein Annexin V,^27^ while 90 % replacement was achieved for recombinantly expressed insulin, likely due to incomplete removal of Pro from the expression host.^28^

Next, we calculated the redox potential of the Trx1Thp variant after determining the disulfide exchange equilibria of Trx1P and Trx1Thp with wild type Trx (E^0^′ = −270 mV) as a reference (Supplementary Figure 2).^18^ Interestingly, the calculated redox potential for Trx1Thp (E^0^ ‘= -266 mV) was more reducing than that of the parent protein Trx1P (E^0^ ‘= -236 mV) by 30 mV and similar to that of wild type Trx. In comparison, all fluorinated Trx1P variants containing (2*S*, 4*S*)-fluoroproline, (2*S*, 4*R*)-fluoroproline, or 4,4-difluoroproline, that we have previously characterised, displayed a redox potential almost identical to Trx1P and were ∼40 mV more oxidising than wild type Trx.^18, 40^

The biological activity of Trx1Thp was assessed by testing its suitability as substrate for bacterial thioredoxin reductase (TrxR), by following the reduction of 5,5-dithio-bis-(2-nitrobenzoic acid) (DTNB) at 412 nm as previously described (**Figure 2a**).^41^

**Figure 2.**
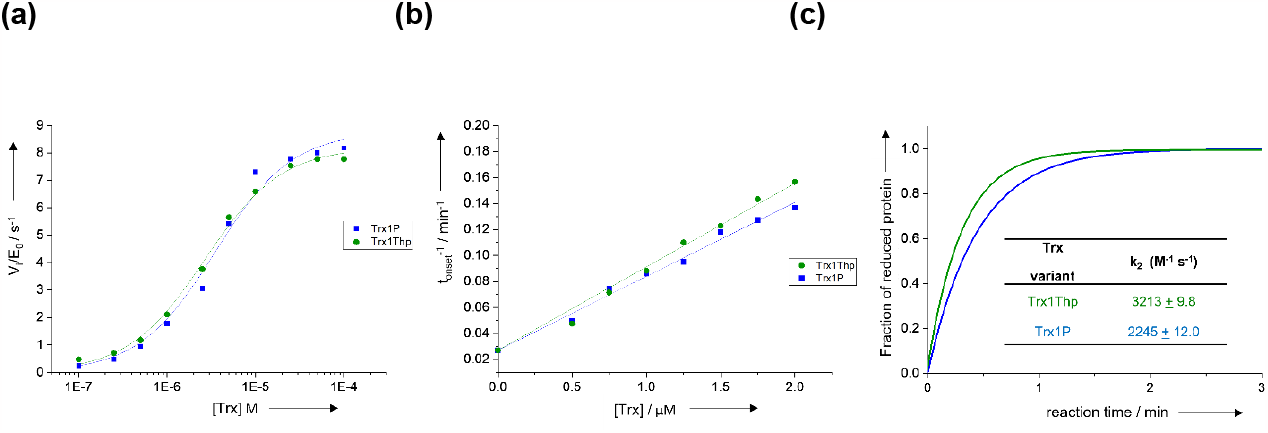
(a) Kinetic analyses of Trx1Thp as substrate of thioredoxin reductase (TrxR) in 50 mM MOPS·NaOH at pH 7.0 and 25 °C in comparison to the parent protein Trx1P (kinetics parameters for Trx1P previously determined under identical conditions are reported from Ref40). (b) Reactivity of the Trx1Thp variant as reductant of non-natural disulfide substrate at pH 7.0 and 25 °C, specifically as catalysts of insulin (0.13 mM) reduction by DTT (1.67 mM) in comparison to the parent protein Trx1P (previously determined under identical conditions, reported from Ref 40). Reactions were followed by the increase in optical density at 650 nm, caused by aggregation of the reduced insulin B chain. (c) Reactivities of the active-site cysteine pairs in Trx1Thp compared to Trx1P (determined previously under identical conditions, reported from Ref40) determined by following the increase in fluorescence at 350 nm (*λ*ex = 280 nm) in the presence of DTTred at pH 7.0 and 25 °C (solid lines). Data are an average of three independent measurements.

The reduction of DTNB proceeds as oxidised Trx (Trx_ox_) is reduced by TrxR, the only enzyme that can reduce Trx_ox_, in the presence of NADPH as reducing equivalent donor. The variant Trx1Thp showed a slightly increased affinity for TrxR in comparison to the parent protein Trx1P (K_M_ = 2.5 ± 0.24 x10 ^-6^ M and 3.0 ± 0.16 ×10^−6^ M respectively) with a similar catalytic activity. These results suggest that the distinct stereochemical and stereoelectronic properties of Thp introduced at position 76 did not result in significant changes in the local tertiary structure and therefore did not affect the interaction between Trx and TrxR. The reductase activity of reduced Trx1Thp was further investigated using bovine insulin as a disulfide containing substrate.^42^ The reaction was monitored by following the increase in optical density at OD_650_ nm, due to the aggregation of the reduced insulin B chains. The aggregation onset was defined as the time taken for the OD_650_ nm to reach 0.05. By plotting the inverse of aggregation onset against the Trx variant concentration, linear plots can be obtained and the insulin reductase activity of each Trx variant can be determined by the slope of the linear regressions (**Figure 2b**). The reductase activity of the Trx1Thp variant was almost identical to that of Trx1P (6.43 ± 0.14 × 10^−2^ min^-1^ µM^-1^ and 5.68 ± 0.11 × 10^−2^ min^-1^ µM^-1^ respectively). Next, we determined the rate of reduction of the disulfide bond active site in the Trx1Thp variant by following the increase in the fluorescence emission at 350 nm (λ_ex_ = 280 nm) upon reduction of the catalytic disulfides with DTT. The rate of reduction for Trx1Thp was 1.4 fold higher than Trx1P (k_2_ = 3213 ± 9.8 M^-1^ s^-1^ and 2245 ± 12 M^-1^ s^-1^ respectively) (**Figure 2c**). These results show that the Pro to Thp replacement at position 76, in proximity of the active site, had no impact on the catalytic activity or the reactivity of the active-site disulfide bonds.

Next we determined the thermodynamic stability of the oxidised and reduced forms of the Trx1Thp variant, by measuring the unfolding/refolding equilibrium transitions at different guanidinium chloride (GdmCl) concentrations (**Figure 3**).

**Figure 3.**
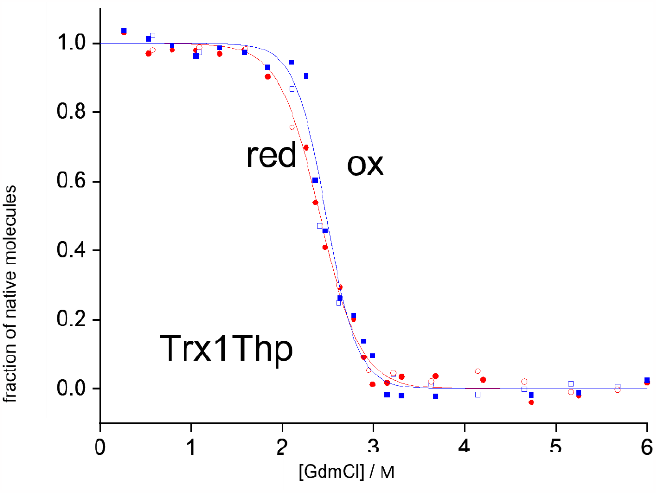
GdmCl-dependent unfolding–refolding equilibria of the oxidised (blue) and reduced (red) forms of Trx1ThpP in 50 mM MOPS·NaOH at pH 7.0 and 25 °C, determined by following the signal change at 220 nm by CD. The unfolding (closed symbols) and refolding (open symbols) equilibria were evaluated according to the two-state model of folding and normalized (solid lines). Data are an average of three independent measurements.

The free energy of folding (ΔG^0^) for the oxidised form (Trx1Thp_ox_) and for the reduced form (Trx1Thp_red_) of the variant were −35.6±1.5 kJ/mol and −26.7±1.8 kJ/mol respectively and were almost identical within experimental error with the previously reported free energy of folding values for the parent protein Trx1P (Table1).^40^ The replacement of the five Pro residues to Thp in annexin V resulted in a decrease in the melting temperature by 4.5 K, although the X-ray structure of the Annexin V variant revealed no effect of the mutations on the overall three-dimensional fold of the protein.^27^ The slight decrease in the melting temperature was therefore attributed to the increased local flexibility of the peptide bond due to the enhanced *cis/trans* isomerisation rate, as well as to solvation effects caused by the increased hydrophilic character of Thp compared to Pro. In fact, all five proline residues in annexin V are solvent exposed, therefore the increased hydrophilicity of the solvent accessible Thp residues may be at least partially accountable for the decrease in stability. However, Pro76 in Trx1P is buried inside the tertiary structure, therefore solvation effects should not play a major role.

The effect of Thp at position 76 on the kinetics of prolyl *cis/trans* isomerization in the context of the intact tertiary structure and in the unfolded state of Trx1P was investigated next. We have previously shown that ∼95% of Trx1P molecules display a *trans*Pro76 in the unfolded state (U^trans^).^41^ These collapse during refolding into a long-lived intermediate with a native-like structure that also features a non-native *trans* Pro76 (I^trans^) in the protein interior. The slow conversion (t_1/2_ = ∼120 min) from I^trans^ to the native state displaying the correct *cis* peptide configuration (N^cis^) represents the bottleneck of the folding reaction (Scheme 1).

**Scheme 1.**
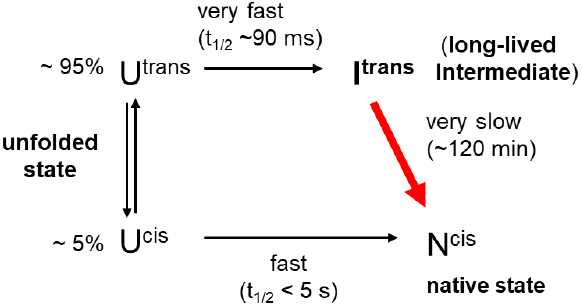
Simplified refolding pathway of Trx1P (adapted from Roderer et al).^41^

Unfortunately, interrupted refolding studies (N-tests) could not be performed for Trx1Thp to assess the fractions of U^cis^ and U^trans^ molecules in the unfolded state and the folding kinetics of the I^trans^ to N^cis^ reaction. In fact, the unfolding reaction in 2.9 M GdmCl (the minimum GdmCl concentration required to unfold >90% Trx1Thp molecules as determined by the equilibrium unfolding transition experiments) proceeded too quickly to be monitored with manual mixing by far-UV circular dichroism. To circumvent this problem, the folding kinetics of I^trans^ to N^cis^ was instead monitored by following the decrease in fluorescence emission at 345 nm (λ_ex_ = 280 nm), as previously described, as I^trans^ displays a twofold higher fluorescence emission intensity at 345 nm than N^cis^.^24, 40^ Briefly, the protein was unfolded overnight in GdmCl (4.0 M) at 25 °C to attain the U^cis^/U^trans^ equilibrium, and subsequently refolded at 25 °C by rapid dilution with MOPS·NaOH (50 mM, pH 7.0) to a final GdmCl concentration of 0.2 M. The rate of the I^trans^ to N^cis^ reaction for Trx1P has been previously determined to be 9.28 ± 0.5 × 10^−5^ s^-1^ with a half-life of ∼120 min.^40^ The replacement of *cis*Pro76 with Thp elicited a 2 orders of magnitude acceleration (>200 fold) of the refolding rate (2.13 ± 0.1 ×10^−2^ s^-1^) with a calculated half-life of 34.5 s (**Figure 4a, Table 2**).

**Table 1.**
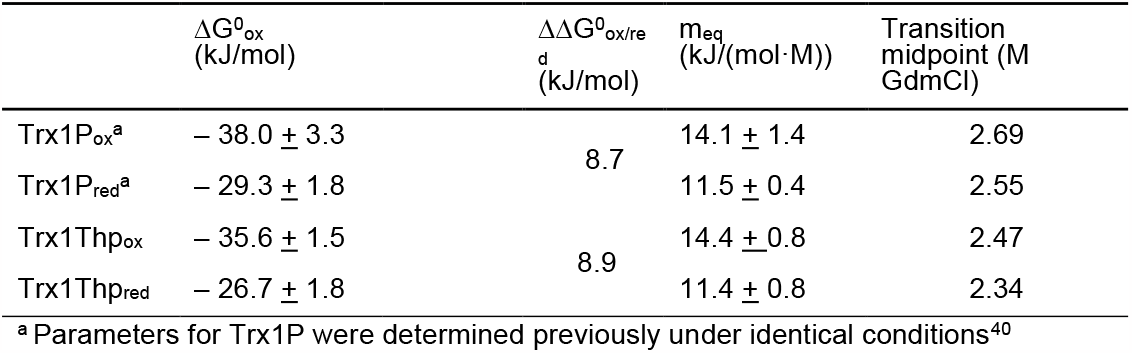
Thermodynamic parameters for the Trx variants determined by unfolding/refolding equilibrium transitions in GdmCl (pH 7.0, 25 °C)

**Table 2.**
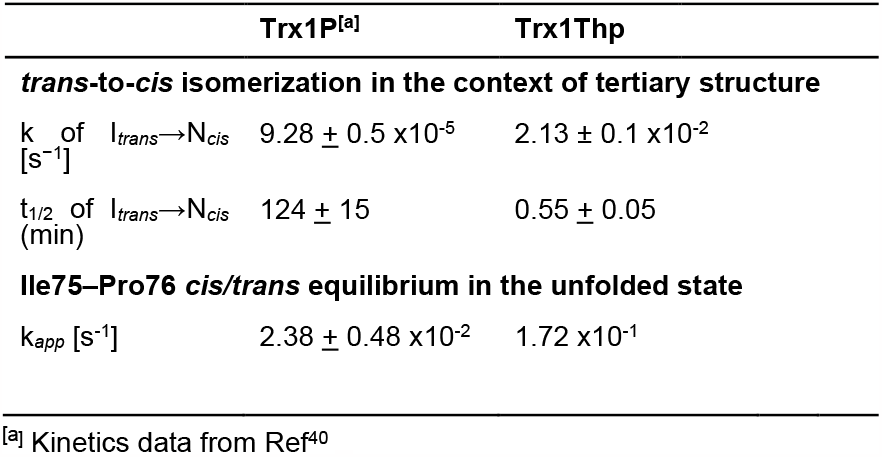
Kinetic folding parameters of Trx1P and Trx1Thp

**Figure 4.**
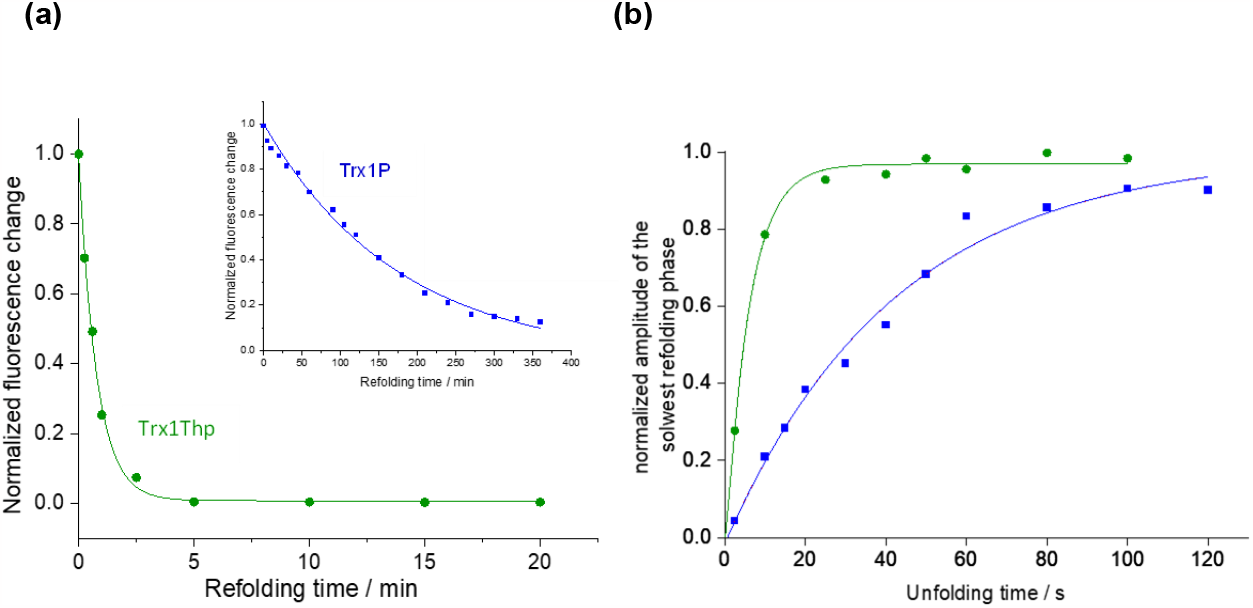
(a) Kinetics of the rate-limiting I^trans^-to-N^cis^ reaction of Trx1Thp, determined by the decrease in tryptophan fluorescence emission at 345 nm, upon excitation at 280 nm. The data were fitted mono-exponentially and normalized. Inset: Kinetics of the rate-limiting I^trans^-to-N^cis^ reaction of Trx1P (previously determined under identical conditions, reported from Ref^40^) (b) Kinetics of the attainment of the *cis/trans* equilibrium of the 75–76 peptide bond after rapid unfolding of the Trx variant Trx1Thp with GdmCl, as determined by interrupted unfolding experiments. After different incubation times in 5 M GdmCl, refolding was measured as the decrease in tryptophan fluorescence at 345 nm (*λ*ex = 280 nm). Kinetics data for Trx1P were previously determined under identical conditions and are reported from Ref ^40^.

As a comparison, the incorporation of (2*S*, 4*S*)-fluoroproline at *cis*Pro76 led to a 9-fold acceleration of the I^trans^ to N^cis^ reaction, while (2*S*, 4*R*)-fluoroproline and 4,4,-difluoroproline had no effect on the refolding rate of Trx1P.^24, 40^ We suggested that a possible explanation for this behaviour might correlate with the pucker preferences of the fluorinated prolines. In fact, only (2*S*, 4*S*)-fluoroproline favours the adoption of the *endo* pucker for which the thioredoxin fold, shows an almost unique preference for the conserved *cis*Pro76 in the context of the tertiary structure. The pseudoproline Thp also displays a preference for the *endo* pucker, therefore we can speculate that this can play at least partially a role in the increase of the refolding rate of Trx1Thp. We have previously reported an acceleration of 4 orders of magnitude in the folding of Trx0P upon Pro76 to Ala substitution in Trx1P.^41^ This mutation generated the formation of a *trans* alanine bond, thus eliminating the slow isomerisation reaction, resulting in a half-life of 90 ms for the formation of native molecules during refolding. Even in the absence of a X-ray structure for the Trx1ThP variant, we are certain that the protein retains the *cis* peptide bond at position 76, as the biological activity and thermodynamic stability of the mutant are almost identical to that of the parent protein. In fact, the small structural rearrangements around the active site of Trx0P resulting from the presence of the *trans*Ala76 bond both impaired the enzymatic activity of thioredoxin and caused a ∼20 kJ/mol loss in the thermodynamic stability of the oxidized form, while the reduced form was 7.6 ± 0.4 kJ mol^−1^ more stable than the oxidized form.

The *cis/trans* equilibrium around the 75-76 peptide bond in the unfolded state was determined by interrupted refolding experiments (U-tests). Briefly, the native protein Trx1Thp was rapidly unfolded in GdmCl (5.0 M) at 25 °C and at different incubation times it was refolded by rapid dilution with refolding buffer to a final GdmCl concentration of 0.1 M. The refolding reactions were recorded as the decrease in fluorescence at 345 nm (λ_ex_ = 280 nm) and the refolding amplitudes were plotted against unfolding time to yield the apparent rate constant (k_app_) of the attainment of the U^cis^/U^trans^ equilibrium (**Figure 4b, Table 2**).

The apparent rate constant k_app_ was determined to be 1.72 ×10^−1^ s^-1^ for Trx1Thp, that is 7-fold faster than the apparent rate for Trx1P (k_app_ = 2.38 + 0.48 ×10^−2^ s^-1^). The microscopic rates constants k_trans→cis_ and k_cis→trans_ (k_trans→cis_ + k_cis→trans_ = k_app_) could not be calculated, as the U^cis^ and U^trans^ fractions in the unfolded state could not be determined by N-tests. Mass spectrometry analysis was performed again after the kinetic measurements. No oxidation of the sulfur of the Thp ring was detected during the storage time. In fact, the diastereomeric S-oxides display different conformational preferences than Thp that might affect the isomerisation rates.^35^

To assess whether the impact of Thp on the acceleration of the I^trans^ to N^cis^ reaction rate observed in the context of the Trx fold, would apply also to other proteins containing *cis* Pro residues in the native state, Thp was incorporated into the model protein pseudo wt barstar (Pro27Ala; Cys40Ala; Cys82Ala) in place of *cis*Pro48. The *in vitro* refolding of pwt-b* has been previously investigated and the fraction of U^cis^ molecules in the unfolded state have been determined to be ∼30 %.^43^ Similarly to Trx1P, the rate-limiting folding step is the isomerization of the peptide bond around Tyr47-Pro48 from the *trans* conformation in the native-like intermediate to the *cis* conformation in the native state. However, while in Trx1P, the non-native *trans* Pro in the structured intermediate is buried in the protein interior, in pwt-b*, Pro48 is located at the surface and is largely solvent exposed.

The incorporation yield for Thp into pwt-b* at position 48 was ∼75% as determined by mass spectrometry (Supplementary Figure S3). Unfolding/refolding transitions in different concentrations urea were performed for the variant pwt-b*-Thp and the parent protein (**Figure 5**). The thermodynamic stability of the pwt-b*-Thp variant was almost identical to that of the parent protein (ΔG^0^ = 14.93 + 1.5 kJ/mol and 14.31 + 3.4 kJ/mol, respectively), while the m_eq_ value of the created variant was higher (7.73 + 0.7 kJ/mol·M) in comparison to pwt-b* (5.29 + 1.3 kJ/mol·M), thus indicating an increase in the folding cooperativity.

**Figure 5.**
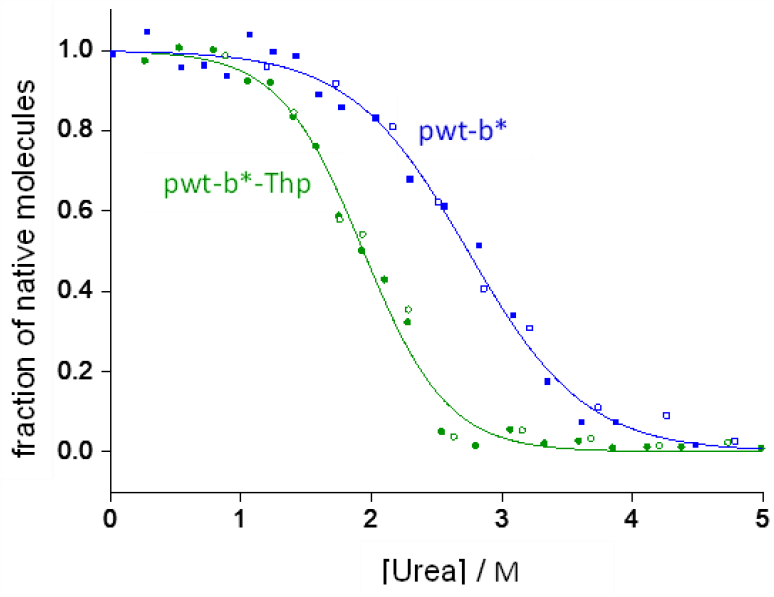
Urea-dependent unfolding/refolding equilibria of pwt-b* (blue) and pwt-b*-Thp (green) at pH 8.0 and 25 °C. Unfolding (closed symbols) and refolding (open symbols) reactions were incubated for 24 h prior to the recording of the far-UV CD signal at 220 nm. Data for each unfolding/refolding equilibrium were fitted globally according to the two-state model of folding and normalized (solid lines).

The proteins pwt-b* and the variant pwt-b*-Thp were fully unfolded in 4 M urea and then refolded by rapid dilution using sodium phosphate buffer. The refolding reaction was followed by the increase in fluorescence emission at 320 nm. The rate of the slow refolding phase (*trans* to *cis* isomerisation around the Tyr-Pro peptide bond) was determined to be 8.2 × 10^−3^ s^-1^ for pwt-b*, in agreement with previously published data. ^43^ As observed for Trx1Thp, the refolding of pwt-b*-Thp occurred much faster than the parent protein. In fact, the refolding reaction occurred within few seconds, and it is therefore too fast to be accurately measured by manual mixing (**Figure 6**).

**Figure 6.**
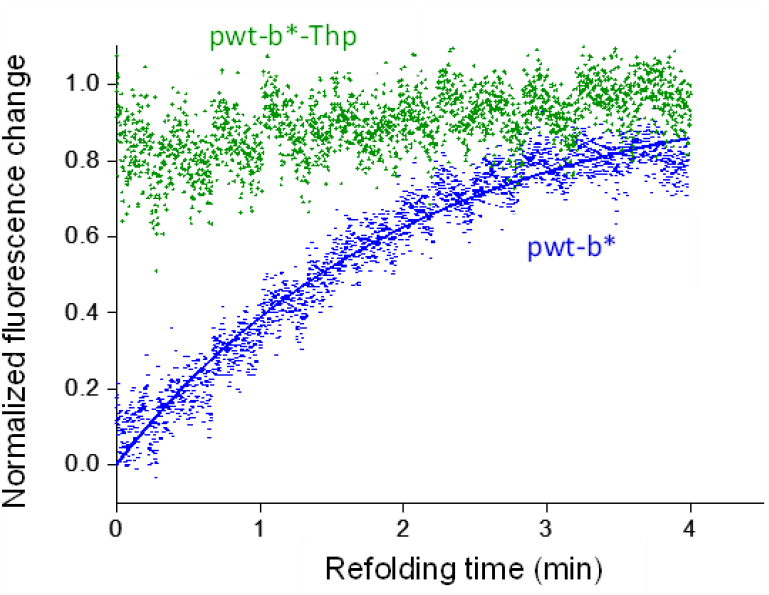
Kinetics of the slow refolding phase of pwt-b* (blue) and pwt-b*-Thp (green), determined by the increase in tryptophan fluorescence at 320 nm, upon excitation at 280 nm. The data for pwt-b* were fitted mono-exponentially and normalized.

Our results suggest that the acceleration of the refolding rate elicited by the presence of Thp might be valid for all proteins containing *cis* Pro residues in their native state. It can be reasoned that the decrease in the activation energy barrier for the *cis/trans* isomerization reaction, together with the preferred adoption of the *endo* pucker and the stabilisation of the *cis* peptide bond all contribute to the dramatic acceleration of the refolding reaction. Early computational studies for the Ac-Thp-NHMe and Ac-Pro-NHMe model peptides indeed indicate that with Thp there is a higher proportion of *cis* conformations.^44^ This was confirmed by experimental studies on model peptides exploiting the *trans*-isomer specific hydrolysis by chymotrypsin. Further, it could be shown that the free energy barrier of isomerisation for a Thp containing peptide is significantly smaller for the *trans*-to-*cis* conversion than for the equivalent Pro containing peptide (ΔΔ_‡_*G* > 20 kJ/mol).^45^ Such barriers will, however, be influenced by the specific peptide context (e.g. hydrogen bonding, steric constraints). Here, we investigated whether the intrinsic chemical difference between Thp and Pro is sufficient to explain the increase in the acceleration of the rate constants for the attainment of the U^cis^/U^trans^ equilibrium in the unfolded state and the I^trans^ to N^cis^ folding kinetics in the context of thioredoxin tertiary structure. We estimated the rotational barriers in Pro and Thp by means of Density Functional Theory (DFT) calculations (M06-2X/cc-pVTZ) on two minimalistic model systems, only including the corresponding 5-membered ring and an acetyl group. This approach was taken to isolate the effect of the substituents in the nature of the peptide bond from other effects, such as steric clashes or changes in the strength of non-covalent interactions (e.g. N···H−N hydrogen bonding) due to the substitution. This allowed us to consistently compare activation energies for the two systems in two different conformational states (orientation of the nitrogen lone pair) and both possible rotation directions (**Figure 7**).

**Figure 7.**
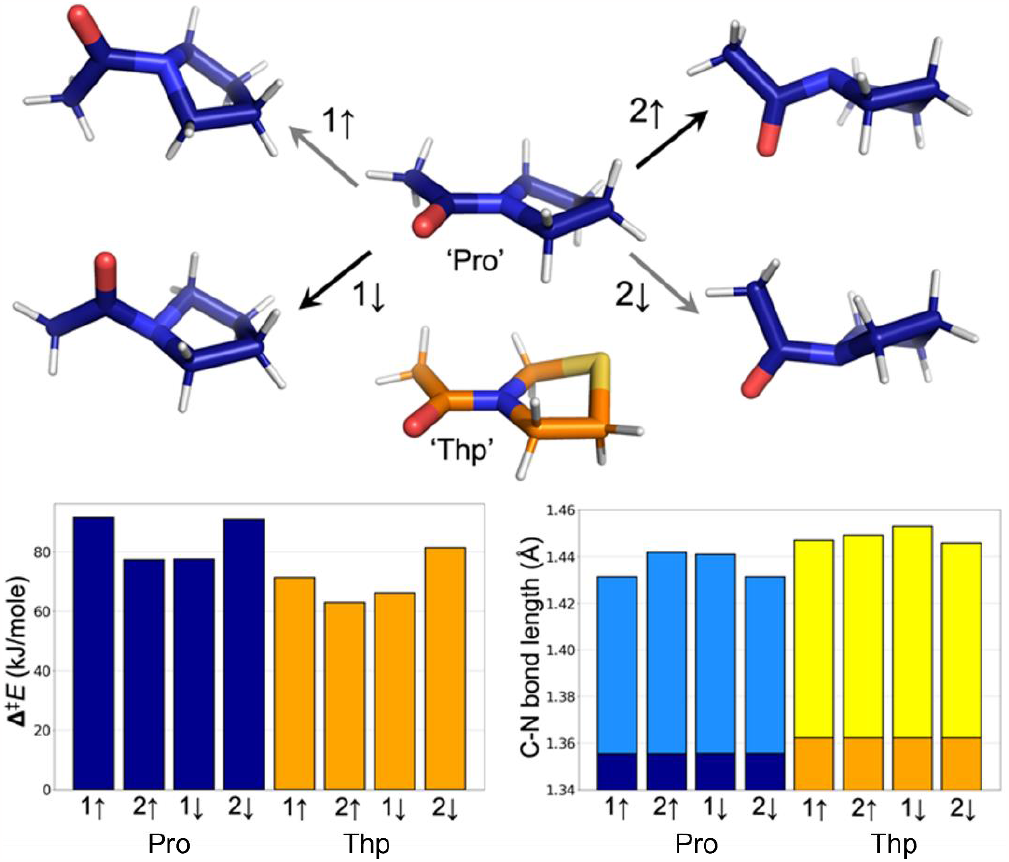
Structures of Pro/Thp minima and Pro transition states, related barrier heights (left bar blot) and C-N bond lengths (right bar plot) for each of the isomerization pathways. C-N bond length bars in dark blue and orange are for the Pro and Thp ground states respectively, while for the transition state the bars are light blue (Pro) and yellow (Thp). The arrows denote the orientation of the nitrogen lone pair and 1 and 2 correspond to the rotation direction.

For the two species, rotation is preferred in the direction that avoids the lone pairs on the oxygen and nitrogen to approach each other (↓1 and ↑2, **Figure 7**). The potential energy barriers for the Thp model are significantly lower than those for Pro model (ΔΔ_‡_*E* > 12 kJ/mol for ↑2), consistent with our experimental results for globular proteins and previous experiments on small peptides,^45^ respectively. In these minimal models, the start- and minimum energy endpoints are structurally indistinguishable, hence there is no notable difference in backward and forward barriers. This indicates that the C^γ^H_2_-to-S replacement in Pro is sufficient to explain the differences in the folding kinetics. The C-N bond lengths in the transition states, a proxy for the double bond character (and thus the level of electron overlap), are in very good agreement with the energy barriers (higher barrier – shorter bond). This trend was not recognized in earlier computational works,^44, 46^ likely due to less accurate quantum mechanical structure optimization (HF/6-31+Gd). These results suggest that the main mechanism by which Thp impacts the isomerization kinetics is by diminishing the double bond character of the peptide bond.

In conclusion, we have shown that incorporation of Thp in place of Pro, accelerates the slow folding phase of both Trx1P and pwt-b* up to 2 orders of magnitude, suggesting that this proline analogue might eliminate the bottleneck of the refolding reaction of proteins containing *cis* proline residues in the native state. Our results also confirmed that Thp behaves as an isostere of Pro, as the incorporation of the analogue was well tolerated by the protein scaffolds which retained their thermodynamic stability and bioactivity. We anticipate that due to its unique features, Thp will play a major role in protein design.

## Materials and Methods

### Proteins Expression and Purification

Trx1Thp was produced in M9 minimal medium using the proline auxotrophic cell strain CAG18515, co-transformed with the plasmids pGDR11-Trx1P and pTARA-ProRS(C443G) similarly as previously described for the variant Trx1Dfp, with the exception that cell growth was performed with 0.035 mM Pro.^40^ Protein expression was allowed to proceed for 3 h at 37 ^°^C in the presence of 8 mM Thp. Protein purification was carried out as described previously.^40^

The variant pwt-b*-Th1P was produced by following the same procedure as above using the CAG18515 *E. coli* strain co-transformed with the plasmid pQE80L–b*(P28A) and pTARA-ProRS(C443G). Protein expression was allowed to proceed for 3 h at 27 ^°^C. The protein was purified as described elsewhere.^43^

### Equilibrium unfolding and refolding transitions

Trx1Thp (17 *μ*M each sample) was incubated at different GdmCl concentrations overnight at 25 °C, in 50 mM MOPS·NaOH, pH 7.0. DTT (5 mM) was added to samples for disulfide bond reduction (Trx1Thp_red_). For measuring refolding equilibrium transitions, the Trx variants were first fully unfolded in buffer containing 6.0 M GdmCl overnight at 25 °C. MOPS·NaOH, pH 7.0 containing different concentrations of GdmCl was then added and samples were then incubated at 25 °C overnight, to allow to attain equilibrium. The CD signal of each sample at 220 nm was recorded for 1 min and averaged, on a JASCO J-810 Circular Dichroism Spectrophotometer. The unfolding and refolding equilibrium of the pwt-b* variants were performed as for Trx1Thp, except urea was used as the chemical denaturant (0.1 – 5 M) in NaH_2_PO_4_, 50 mM, pH 8.0. The average CD signals were then plotted against the GdmCl concentrations. The data were fitted according to the two-state model of folding and normalized (Equation 1).

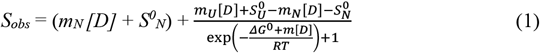

m_n_ and m_u_ are the “slopes” of the pre- and post-transition baselines; [D] is the denaturant concentration; *S*^0^_*n*_ and *S*^0^_*n*_ are the signals of native and unfolded protein at 0 M denaturant; ΔG^0^ is the free energy of folding at 0 M denaturant; and *m* is the linear dependence of ΔG^0^ on [D].

### Refolding Kinetics Measurements

Trx1Thp was unfolded overnight in buffer containing 4.0 M GdmCl at 25 °C to attain the U^cis^/U^trans^ equilibrium, and subsequently refolded at 25 °C by rapid dilution (1:20) with MOPS·NaOH (50 mM, pH 7.0) to a final GdmCl concentration of 0.2 M. The pwt-b* variants were unfolded overnight in buffer containing 5.0 M urea, at 25 ^°^C, and refolded by rapid dilution (1:20) with NaH_2_PO_4_ (50 mM, pH 8.0), to a final urea concentration of 0.25 M. The refolding reaction was followed on a Cary Eclipse Fluorescence Spectrophotometer at 345 nm emission (*λ*_ex_ = 280 nm). The decrease (Trx1Thp) or increase (pwt-b* and pwt-b*-Thp) in fluorescence emission was plotted against reaction time (min) and fitted monoexponentially to give the rate of I^trans^ to N^cis^ conversion.

### Interrupted Unfolding Kinetics Measurements

Native Trx1Thp (200 μM) was rapidly unfolded in buffer (MOPS·NaOH, 50 mM, pH 7.0) containing 5.0 M GdmCl at 25 °C, incubated for different times under these conditions, and then refolded at 25 °C by rapid dilution (1:50) with MOPS·NaOH (50 mM, pH 7.0) to a final GdmCl concentration of 0.1 M. The refolding reactions were recorded as the increase in fluorescence at 345 nm (λ_ex_ = 280 nm), on a Cary Eclipse Fluorescence Spectrophotometer. Refolding kinetics were fitted mono-exponentially with initial and final signal intensity as open parameters. The refolding amplitudes were plotted against unfolding time and yielded the apparent rate constant (k_app_) of the attainment of the U^cis^/U^trans^ equilibrium.

### Trx1Thp as a substrate for bacterial TrxR

Experiments were carried out using a quartz cuvette and a Cary-50 UV-Visible spectrophotometer, at 25 °C, in 50 mM MOPS·NaOH, pH 7.0, 0.4 mM EDTA. Recombinant *E. coli* TrxR was incubated with DTNB. Solutions of various concentrations of Trx1Thp were added to the TrxR/DTNB solution and the reaction was started by addition of NADPH. The final concentrations were as follows; TrxR 33 nM, Trx 0.1–300 µM, DTNB 3.3 mM and NADPH 0.8 mM. TrxR activity was determined by monitoring the change in absorbance at 412 nm during the initial 60 seconds of the reaction. Slopes were converted into initial velocities using the extinction coefficient ε_412nm_ = 28300 M^-1^ cm^-1^, corresponding to the formation of two TNB molecules per catalytic cycle. Initial velocities were plotted against Trx concentrations and data fitted according to Michaelis-Menten kinetics.

### Reduction Kinetics of Trx1Thp variants by DTT

Equilibrium fluorescence spectra of the Trx variants were recorded between 320 and 400 nm, with excitation at 280 nm, using a Cary Eclipse Fluorescence Spectrophotometer. The reduction of the oxidized form of the Trx variants was measured by addition of reduced DTT with a final concentration of 2 µM Trx and 20 nM DTT in 100 mM NaH_2_PO_4_-NaOH, pH 7.0. The sample was excited at 280 nm and the increase in emission was recorded at 350 nm. The monoexponential fitting was used to calculate the rate of reduction.

### Insulin Reduction Assay

The catalytic activity of Trx1Thp on the reduction of bovine insulin by DTTred was measured in 100 mM KH_2_PO_4_, pH 7.0, 2 mM EDTA. Trx1Thp was incubated with DTT_red_ for 5 min and the reaction was started by addition of the bovine insulin. The end concentrations were as follows; DTT_red_: 1.67 mM; bovine insulin: 130 µM and Trx: 0.5, 0.75, 1, 1.25, 1.5, 1.75, 2 µM. The reaction was monitored via the increase in optical density at 650 nm (OD_650nm_). Aggregation onset was measured as the time taken for the optical density at 650 nm to reach 0.05, measuring on a Jenway 7315 Spectrophotometer. The reciprocal of the aggregation onset was plotted against Trx1Thp concentration to obtain linear plots, where the slope of the linear fit was the rate of reaction.

### Density functional theory calculations

The reaction pathways and energy profiles were obtained for the two model systems: 1- (pyrrolidin-1-yl)ethan-1-one and 1-(thiazolidin-3-yl)ethan-1-one. For each system, four isomerization pathways were obtained – one for each combination of the nitrogen state and rotation direction. The same protocol was used for all the pathways. First, the potential energy scan along the peptidic dihedral angle rotation was performed using GFN2-xTB to obtain an initial guess for the initial and final structures as well as the transition structure.^47^ Then, the pathways were optimized using the Climbing-Image Nudged Elastic Band (CI-NEB)^48^ implementation in ORCA v. 5.0.1.^49^ The M06-2X^50^ functional with the cc-pVTZ basis set^51^ was employed. The protocol includes preliminary optimization of the two end-point structures to ensure that they correspond to minima on the potential energy surface.

## Acknowledgements

Marina Rubini and Jennie O’ Loughlin are grateful to the UCD School of Chemistry and to the Seed Funding Scheme (SF1888) for financial support. Marina Rubini would like to thank Vincent Conticello for the pWK2 plasmid containing the *E. coli* ProRS (C443G) gene. Marc van der Kamp and Kirill Zinovjev thank BBSRC for funding (BB/R001332/1).

## Notes

### Competing Interest Statement

The authors have declared no competing interest.

