## Supplementary Figures for "4-Thiaproline accelerates the slow folding phase of proteins containing *cis* prolines in the native state by two orders of magnitude"

**1. Materials**

**1.1 Determination of the redox potential of Trx1Thp**

**2. Supplementary Figures**

**3. References**

### 1. Materials

The following were obtained from Sigma: NADPH tetrasodium salt, 5,5'-dithiobis-2-nitrobenzoic acid (DTNB), insulin from bovine pancreas. L-Thiazolidine-4-carboxylic acid (98% purity) was purchased from Alfa Aesar. *E. coli* TrxR was a gift from Prof. Rudi Glockshuber (ETH Zurich).

#### 1.1 Determination of the redox potential of Trx1Thp

Redox potentials were determined via their equilibrium constant, using Trx wt as a reference, following published procedures.[1] First, Trx wt was freshly reduced in 1 mM DTT and subsequently all Trx variants were buffer exchanged against 50 mM MOPS·NaOH pH 7.0, 1 mM EDTA using PD MiniTrap G-25 columns (GE Healthcare) (500 µL protein load, 200 µL wash and 700 µL elution volume). Oxidized Trx variants were mixed with reduced Trx wt in a final volume of 100 µL (Trx variant 10 µM, Trx wt 10-20 µM), and incubated for 17 hours at 25°C to attain equilibrium. The disulfide exchange reaction was quenched by addition of 12 µL formic acid and 50 µL were loaded on a ZORBAX C8 reverse phase column (300 Å, 4.6 x 250 mm, from Agilent), equilibrated with 35% acetonitrile in water, 0.1% TFA. Protein were eluted with a gradient from 35 to 65% acetonitrile in water, 0.1% TFA over 30 min, with a flow rate of 1 mL/min and a temperature of 70°C. Elution profiles were recorded following the absorbance signal at 220 nm, and protein concentrations calculated via their peak area (using the Peak Analyzer tool of OriginPro 2018b, OriginLab). Redox equilibrium constants were calculated according to Equation 1 and obtained $K_{eq}$ were used to determine the redox potential ($E'_0$) of each Trx variant, according to Nernst equation (Equation 2), using $E'_0$ of the Trx $wt_{red}$/Trx $wt_{ox}$ (-270 mV) redox couple as a reference.[2]

Equation 1: $K_{eq} = \frac{[Trx\ variant_{red}][Trx\ wt_{ox}]}{[Trx\ variant_{ox}][Trx\ wt_{red}]}$

Equation 2: $E'_0 = -270\ mV + (\frac{RT}{2F} lnK_{eq})$

### 2. Supplementary Figures

#### Supplementary figure 1

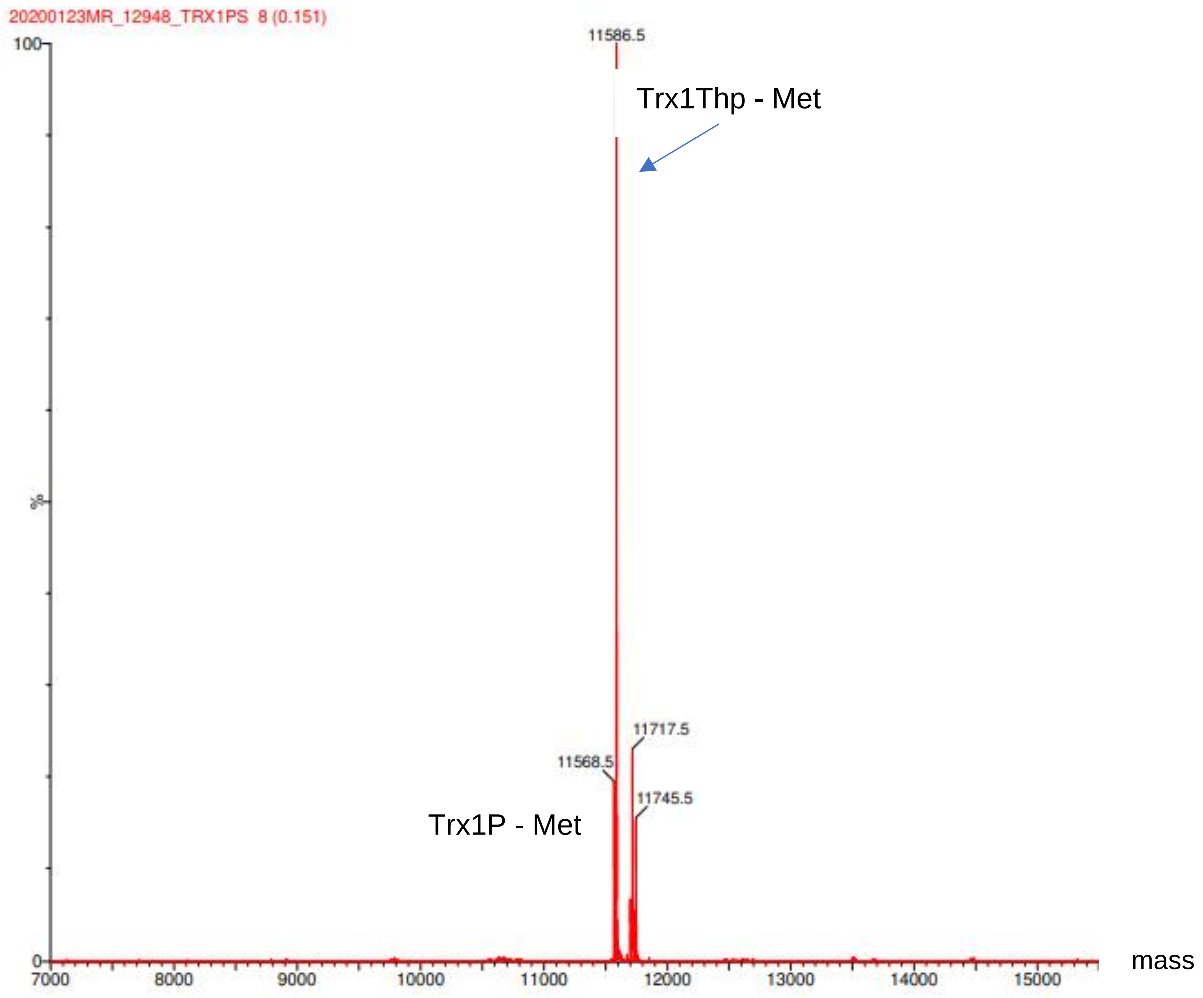


**Supplementary Figure 1.** Deconvoluted ESI-MS spectrum of *E. coli* Thioredoxin variant Trx1Thp.

### Supplementary figure 2

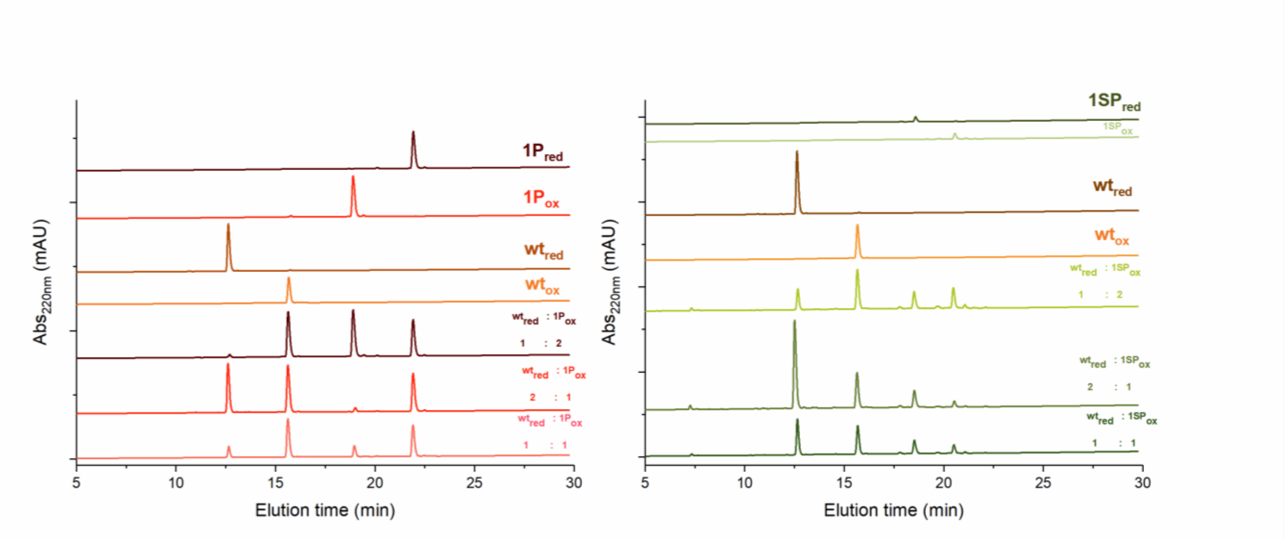


**Supplementary Figure 2. Redox equilibria between Trx wt and Trx1P or Trx1Thp (1SP) at pH 7.0 and 25°C.** Reduced Trx wt was mixed with the oxidized form of the respective Trx variant at a 1:1, 1:2 or 2:1 molar ratio. The reactions were quenched with formic acid, and all redox forms separated by reversed phase HPLC. Peak areas were converted to concentrations, from which redox equilibrium constants ($K_{eq}$) were calculated. $K_{eq}$ values were found to be independent of the mixing ratio between reduced Trx wt and oxidized Trx1P or Trx1Thp, showing that the redox equilibria were attained.

### Supplementary figure 3

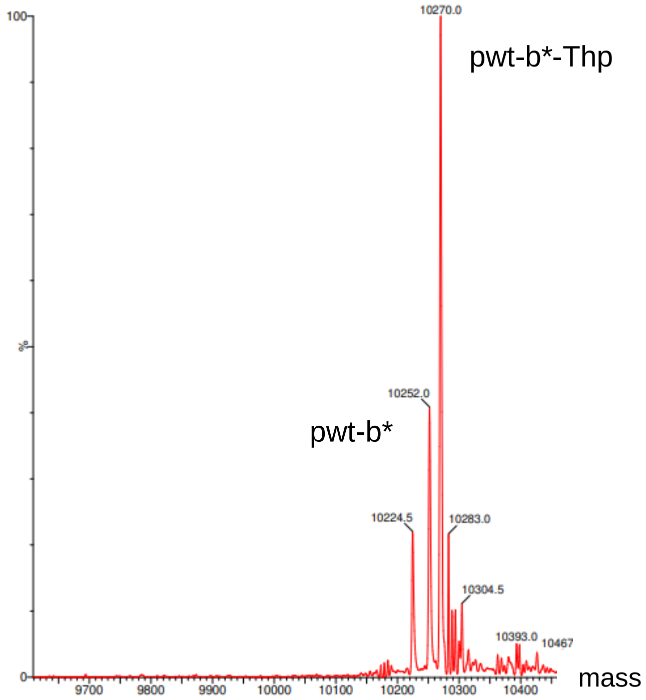


**Supplementary Figure 3.** Deconvoluted ESI-MS spectrum of *B. amiloliquefaciensi* barstar variant pwt-b*-Thp (calculated: 10270.6 Da; found: 10270.0 Da).
